# MitoDate: a Nextflow pipeline for molecular clock dating and phylogenetic inference using ancient mitogenomes

**DOI:** 10.64898/2026.07.28.741234

**Authors:** Wenxi Li, Bilal Sharif, Peter D. Heintzman, Love Dalén, J. Camilo Chacón-Duque

**Author notes:** These authors contributed equally (Wenxi Li, Bilal Sharif).

## Abstract

**Summary:** Ancient DNA studies are increasingly targeting samples that are beyond the limit of radiocarbon dating (>50 thousand years old) and are often difficult or impossible to date using other geochronological methods. In cases where complete mitochondrial genomes (mitogenomes) can be recovered from such samples, Bayesian molecular clock dating approaches are routinely used as an alternative method for estimating their age. However, molecular clock dating of ancient mitogenomes lacks a standardised, reproducible computational framework, and existing approaches rely heavily on graphical interfaces that limit automation and scalability. To address these gaps, we developed MitoDate, an automated Nextflow pipeline for reproducible molecular clock dating of ancient mitochondrial genomes. The workflow standardises Bayesian time-calibrated phylogenetic inference within a portable, containerised framework, reducing manual intervention and improving analytical consistency.

**Availability and implementation:** MitoDate is implemented in Nextflow and is freely available at https://github.com/CpgSthlm/MitoDate. The pipeline is distributed with containerised dependencies and detailed documentation, including example datasets and usage guidelines.

## 1. Introduction

The production of palaeogenomic datasets has increased sharply over the past decade, driven both by continuous improvements in molecular techniques to work with ancient DNA (aDNA), and by the long-term decrease in DNA sequencing costs. As a consequence, the temporal range at which ancient DNA data can be retrieved has also considerably increased into the deep past (Dalén *et al*. 2023). One main challenge to contextualise and analyse palaeogenomic data is to confidently establish the age of the specimens from which the DNA data is obtained. The most widely used approach, radiocarbon dating, has a temporal limit of ∼50,000 years (Reimer *et al*. 2020), meaning that any specimen older than this limit requires a different dating strategy. Other geochronological dating methods can be applied, but none are broadly applicable and either are only useful in specific contexts (*e*.*g*., Uranium series, tephra, optically stimulated luminescence (OSL)), require specimens to be found in situ (*e*.*g*., paleomagnetism, OSL, tephra), offer ages that require local calibration datasets (*e*.*g*., amino acid racemization), or, when used in combination, provide bracketed age information only (Dalén *et al*. 2023). Indirect approaches such as archaeological context and/or stratigraphic information are also often employed, but these typically provide very broad time ranges and are difficult to compare across studies. One widely used alternative is the use of genetic information to perform molecular clock dating.

When sufficient DNA is preserved, complete mitochondrial genomes (mitogenomes) can often be reconstructed, making them one of the most commonly generated genomic resources in ancient DNA studies and a widely used source of phylogenetic information (Paijmans, Gilbert, and Hofreiter 2013). Mitogenomes are also particularly suitable for molecular clock dating, especially for remains that are beyond the effective range of radiocarbon dating, are not amenable to other direct dating methods, or lack reliable contextual age information (Shapiro *et al*. 2011). Molecular clock dating, often performed within a Bayesian framework, estimates evolutionary rates by linking genetic divergence to time through calibration priors based on fossil-calibrated divergence times or previously estimated mutation rates, enabling the inference of both divergence times (node dating) and sample ages (tip-dating). These time-calibrated phylogenetic methods also incorporate known ages of ancient sequences, usually obtained through radiocarbon dating, and account for branch shortening and missing substitutions along unsampled lineages. Bayesian tools such as BEAST (Suchard *et al*. 2018) have therefore become widely used for molecular tip dating analyses of ancient mitogenomes (Rey-Iglesia *et al*. 2019, Karpinski *et al*. 2020, Baca *et al*. 2023, Chacón-Duque *et al*. 2025, Salis *et al*. 2025, Walker *et al*. 2025).

Despite its broad application, molecular clock dating of ancient mitogenomes remains difficult to perform in a standardised and reproducible manner. A typical analysis requires a set of dataset-specific decisions, including the selection of time-calibrated reference samples, specification of tip-dating priors, root calibration, substitution model, molecular clock model, population model, and Markov chain Monte Carlo (MCMC) settings. These settings can directly influence age estimates, yet they are often specified through Graphical User Interfaces (GUI) such as BEAUti (Suchard *et al*. 2018) and study-specific scripts built around fixed templates (Rieux and Khatchikian 2017, BEASTGen, BEAST Documentation, n.d.). In particular, template-based approaches may limit flexibility when datasets require different root calibration strategies, model settings, sample-specific analyses, or iterative re-runs. Recent studies further suggest that independent single-sample dating provides more reliable estimates than jointly analysing multiple undated samples (Karpinski *et al*. 2020, Chacón-Duque *et al*. 2025). While improving dating accuracy, the single-sample dating strategy substantially increases the number of individual BEAST analyses that must be configured, executed, and assessed for convergence. As a result, molecular tip dating workflows often remain difficult to scale, compare, and reproduce, especially when analyses involve multiple undated samples that require independent inference, convergence assessment, and selective reanalysis.

To address these challenges, we present MitoDate, a modular Nextflow pipeline (Di Tommaso *et al*. 2017) for Bayesian molecular tip dating of ancient mitogenomes using BEAST1, building upon the methodology implemented in Chacón-Duque *et al*. (2025). MitoDate standardises and automates dataset preparation, configuration-driven BEAST1 XML generation, Bayesian tip-dating inference, summarisation of results, and joint phylogenetic analyses within a portable and containerised workflow. The pipeline also supports selected-sample reruns, allowing users to confidently estimate sample ages by inspecting convergence and incorporating reviewed estimates into the joint phylogeny. We reproduce the molecular clock dating analyses performed in Chacón-Duque *et al*. (2025) and demonstrate that it can be achieved while reducing manual intervention and improving reproducibility.

## 2. Pipeline implementation

MitoDate is implemented as a modular Nextflow pipeline and features three distinct run modes/workflows (Fig. 1). The first mode, “*tip_dating”*, processes the input multiple sequence alignment (MSA), generates BEAST-compatible XML files, runs the Bayesian tip-dating analyses, and summarises the resulting posterior estimates into a CSV file. The second mode, “*rerun_samples*”, re-runs the tip-dating on selected samples from a previous run without repeating the full mode 1, which is useful when some samples require longer MCMC sampling or adjusted settings. The third mode, “*joint_tree*”, uses the age estimates produced by the tip-dating workflow (modes 1 or 2) to construct a final time-calibrated phylogeny for the full dataset. Together, these modes support an iterative analysis strategy in which individual sample ages can be estimated, inspected, selectively re-run, and incorporated into a joint phylogenetic inference.

**Fig. 1:**
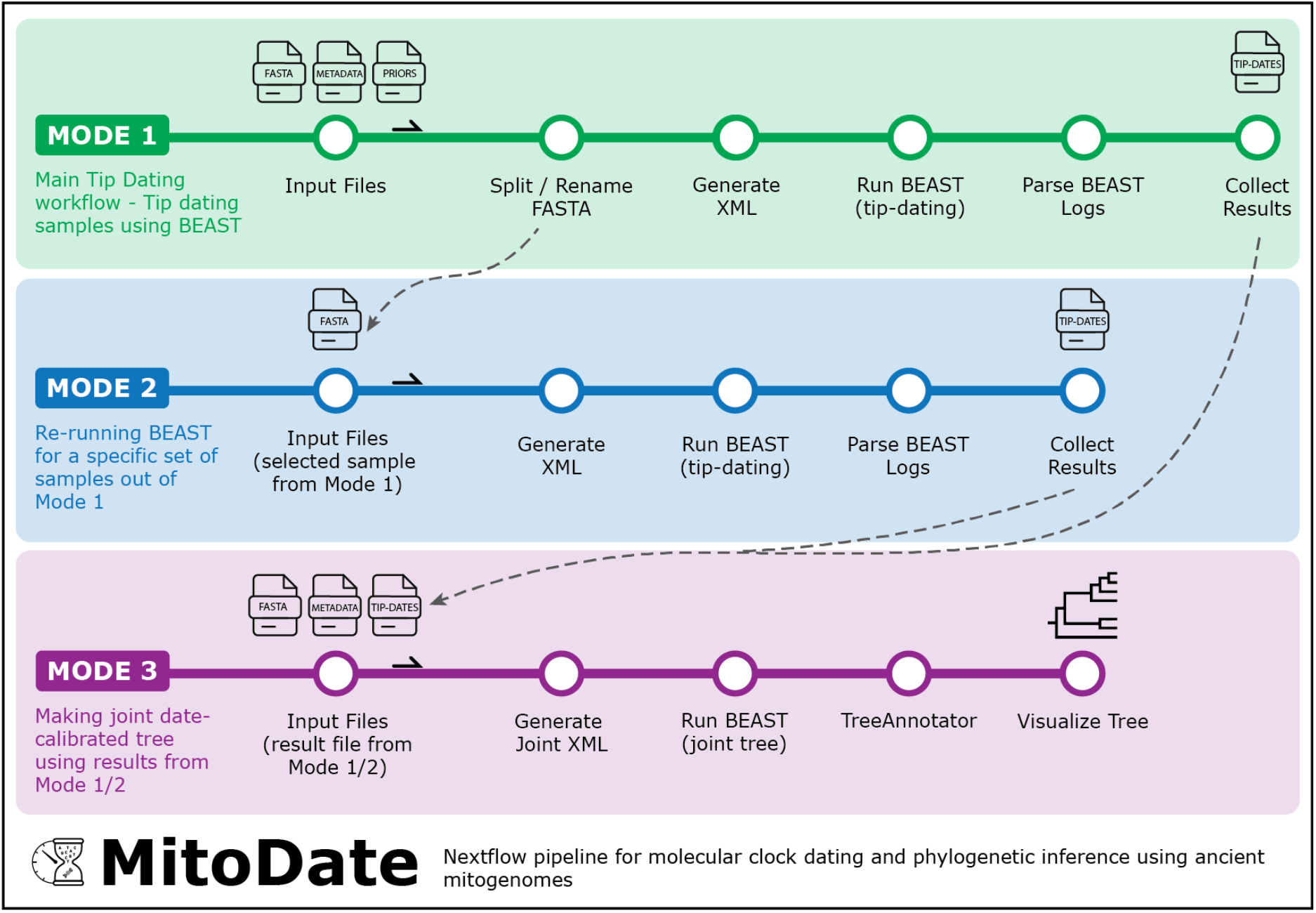
Schematic overview of the MitoDate pipeline. Workflow map of the three run modes: *tip_dating* (Mode 1, green), *rerun_samples* (Mode 2, blue), and *joint_tree* construction (Mode 3, purple).

### 2.1. Data input and configuration

MitoDate is configured using a set of input data files specified in a configuration file. The sequence data is provided as an MSA in FASTA format, in which all sequences must be the same length, and each header corresponds to a sample identifier. A tab-separated (TSV) metadata file then describes each sample, its species, taxonomic or other analytical grouping, its calibrated radiocarbon age point estimate where available, and whether it should be included in the *tip_dating* workflow, while a comma-separated (CSV) priors file provides the age priors used to initialise the analysis for the undated samples. By default, the pipeline does not use partitioning of the mitogenome. If the user prefers to set partitions, an annotation file in GFF format needs to be provided (see Section 2.3 and the online documentation). Because the same sample identifiers are shared across the alignment headers, metadata, and priors file, these inputs are linked throughout the workflow. Before a run begins, MitoDate verifies that the files required for the selected mode are present. All remaining aspects of the analysis, including run mode, model settings, MCMC parameters, execution behaviour, and output directory, are specified in the configuration file.

### 2.2. Sequence processing

In the sequence processing step, the pipeline relabels the input FASTA sequences according to the metadata file and produces either one FASTA alignment per undated sample (single-sample dating mode, the default) or a single combined alignment (multiple-sample dating mode) (see Section 2.4). We highly recommend visually inspecting the MSA before starting the analysis to remove any regions that are unsuitable for reliable molecular tip dating, such as parts of the hypervariable region (*e*.*g*., VNTR) or sites that are only present in one sample and likely represent a sequencing, mitogenome assembly, or MSA generation error.

### 2.3. XML generation

The XML generation module is the central component of the pipeline, which generates the BEAST-compatible XML file directly from the FASTA alignment, metadata, priors table, and configuration parameters, without having to rely on a GUI. Each XML encodes the complete analysis setup, including the taxon labels and tip dates, the substitution and molecular clock models, the population (tree) prior, the root calibration, and the MCMC settings. Parameters not specified in the configuration file retain their BEAST defaults. The tip-date priors are taken from the priors file and may be specified with uniform, normal, or lognormal distributions. When partitioning is enabled, the module assigns the user-specified substitution models to their respective regions of the alignment, delimited by the GFF annotation file, such as protein-coding genes, rRNAs, tRNAs, and the control region. Automating this step removes a common source of manual error and keeps the model specification transparent and consistent across samples.

### 2.4. BEAST tip-dating

The BEAST tip-dating module performs Bayesian inference using BEAST v1.10.4 (Suchard *et al*. 2018) by running the MCMC to sample the posterior distribution of the model parameters and divergence times defined in the XML. Here, MitoDate supports both single-sample and multiple-sample tip-dating strategies, but we strongly encourage the use of single-sample dating (the pipeline’s default). For this strategy, each undated sample is tip-dated using an independent BEAST run, together with all the calibrated radiocarbon-dated sequences. This is recommended as per Chacón-Duque *et al*. (2025), where the authors demonstrated that simultaneously estimating the ages of multiple undated samples within the same tip-dating run can introduce biases, pushing samples to appear older than they are. The multiple-sample approach is retained as an optional mode for users who wish to analyse several target samples simultaneously and compare the results with the single-sample dating estimates. To reduce runtimes, the module supports multi-threaded execution and uses BEAGLE hardware acceleration when available (Suchard and Rambaut 2009).

### 2.5. Result parsing

The result-parsing module extracts and summarises the posterior estimates from the BEAST log files, in a similar way to Tracer (Rambaut *et al*. 2018) (BEAST’s GUI for checking the results). For each analysis, MitoDate outputs the posterior mean, standard deviation, and 95% highest posterior density (HPD) interval for the estimated tip and root ages and the clock rate, together with the effective sample size (ESS) of these and the likelihood terms as a measure of convergence. The per-sample summaries are then aggregated into a single consolidated CSV table, allowing age estimates to be compared across samples. The user can then inspect whether an analysis has converged. At a minimum, we suggest ESS values >200 for both the age estimate and joint probability distribution to assume convergence. Mode 2 *rerun_samples* can be used to re-run specific samples that fail to converge.

### 2.6. Joint tree construction

The mode 3 *joint_tree* re-uses the XML generation and BEAST steps described above to build a final time-calibrated phylogeny, but applies them to the full alignment with all tips treated as dated. It first merges the reviewed age estimates from a previous tip-dating run into the metadata, so that no priors file is required, and then generates and runs a single joint XML for the complete dataset. It is important to note that for the tip-dated samples the pipeline includes both the mean and the 95% HPD values as a measure of uncertainty, while only a point estimate (*e*.*g*., median or mean) for the calibrated radiocarbon-dated samples is included. The resulting posterior tree distribution is summarised into a maximum clade credibility (MCC) tree with TreeAnnotator (TreeAnnotator, BEAST Documentation, n.d.), which MitoDate finally renders as a figure with tips coloured by the user-defined grouping column in the metadata. This mode can also be used with a dataset that contains only radiocarbon-dated and/or present-day samples.

### 2.7. Reproducibility and portability

Reproducibility and portability are central to the design of MitoDate. Workflow execution is managed by Nextflow (Di Tommaso *et al*. 2017), which tracks each process and caches completed tasks for reuse whenever the inputs, parameters, and code are unchanged. Each run can additionally be documented through Nextflow’s execution reports, timelines, and trace files. Together with the text-based configuration file, in which all analytical choices are recorded rather than set through a GUI, this keeps every dating analysis fully traceable and easy to reproduce or share.

Software dependencies are handled through containerised execution, with support for Docker (Merkel 2014), Apptainer/Singularity (Kurtzer *et al*. 2021), and with Conda or Mamba as alternatives. This avoids version conflicts and keeps results portable across local workstations, high-performance computing (HPC) clusters, and cloud environments. For HPC systems, MitoDate ships with predefined execution profiles, and users can add their own by specifying scheduler, CPU, memory, wall-time, and SLURM parameters.

Finally, through Nextflow’s task-based execution model, the individual modules are scheduled as independent tasks and distributed across the available resources. Because the sample-specific analyses are independent, they can run in parallel. This is particularly valuable for molecular tip dating, where computational cost scales with the number of sequences, the alignment length, and the MCMC chain length.

## 3. Application and validation

We evaluated MitoDate’s performance by replicating the tip-dating analysis of the 10 oldest samples from the mammoth mitogenome study of Chacón-Duque *et al*. (2025), ranging in average age from ∼1.4 to ∼0.2 million years ago. This dataset represents a suitable validation case because it contains ancient mitochondrial genomes spanning a wide temporal range. To reproduce the results, we provide the MSA in FASTA format, sample metadata, tip-dating priors, substitution model, clock model, population model, calibration priors, root calibration, and MCMC parameters in accordance with Chacón-Duque’s study. The analysis was then performed using the MitoDate pipeline v0.1.0.

MitoDate successfully reproduced the main molecular tip dating patterns published in the Chacón-Duque *et al*. (2025) analysis (Fig. 2). Across the 10 selected samples, the replicated posterior age estimates were broadly consistent with and almost identical to the published estimates. For most samples, both the means and their 95% HPD intervals showed substantial overlap. For the four oldest samples, the original results seem to be slightly younger, with the six remaining samples displaying almost identical results. It is also important to note that MitoDate’s results with and without partitions are nearly identical, which means that partitioning is likely not necessary when using almost-complete mitogenome sequences. For this reason, we set “no partitioning” as the default for our pipeline.

**Fig. 2:**
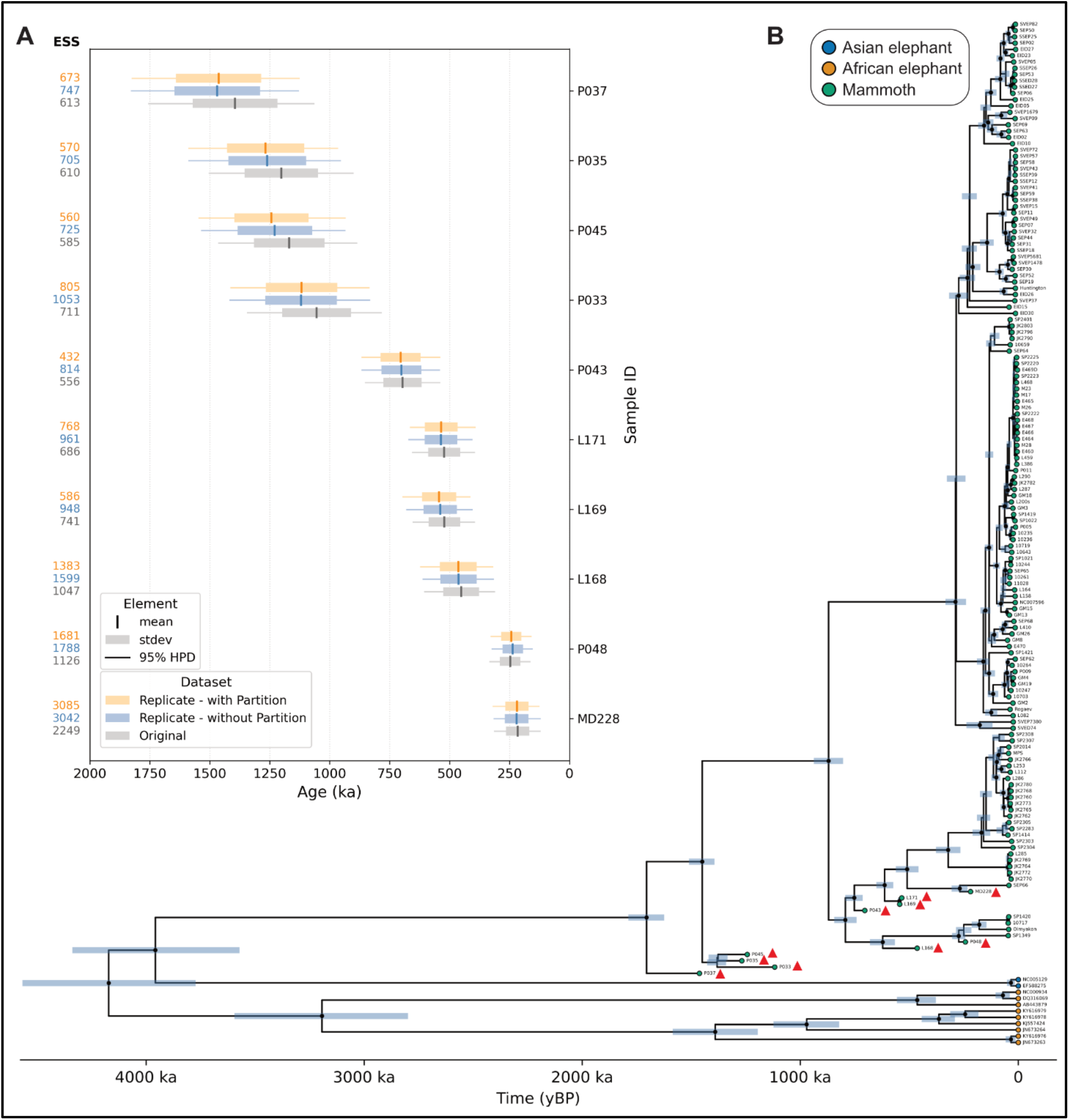
Validation of MitoDate against published age estimates. **(A)** Posterior age estimates from MitoDate compared with the published data of Chacón-Duque *et al*. (2025). (**B)** Joint phylogeny using both single-sample tip dating and radiocarbon estimates as input for tip ages. The blue node bars display the 95% HPD node heights estimated by the Bayesian analyses. Nodes with posterior probability >0.90 are indicated by black dots. Red triangles indicate samples dated using MitoDate single-sample dating.

Together, these results show that MitoDate can reproduce the results from Chacón-Duque *et al*. (2025) under a standardised and automated workflow, overcoming the need for manual configuration and result-processing steps. The validation also demonstrates the practical utility of the pipeline for iterative quality control: posterior means, standard deviations, 95% HPD intervals, and effective sample size values are automatically extracted for each sample, allowing users to evaluate both point estimates and posterior uncertainty, identify analyses that require additional sampling, and re-run selected samples without repeating the full workflow.

## 4. Discussion

MitoDate replaces the manual and template-based steps currently used in molecular clock dating with a single configuration-driven workflow. This makes the assumptions of each dating analysis explicit and allows them to be recorded, shared, and re-run without modification. The pipeline is implemented in Nextflow, which records every process, caches completed tasks, and can reproduce an entire analysis from the same inputs, parameters, and code. Moreover, all of the pipeline’s dependencies are packaged in containerised environments that fix the exact software versions used. Together, this ensures that the same result can be obtained on a local computer, an HPC cluster, or a cloud system, and because each analysis runs as an independent task, the workflow scales well with study size.

Our validation against the mammoth mitogenome dataset of Chacón-Duque *et al*. (2025) confirmed that the pipeline recovers published posterior age estimates. The small differences in the mean ages reflect a methodological refinement in MitoDate, in which we avoid duplicating sequence data where the annotation contains overlap between different partitions, in contrast to the original analysis in Chacón-Duque *et al*. (2025).

Naturally, some challenges and limitations remain. The Bayesian nature of molecular clock dating makes the final estimates highly sensitive to the priors and parameters chosen. For example, relying on a divergence estimate to calibrate the root of the tree inevitably introduces bias, particularly for the oldest samples that are not informed by radiocarbon-dated tips. However, by targeting samples that are well beyond the radiocarbon dating limit but are amenable to alternative geochronological dating methods, it should become possible to constrain the rate of evolutionary change more accurately at deeper points in time. Such improvements could easily be adapted to our pipeline.

Overall, an automated and streamlined workflow for molecular clock dating is a valuable contribution to the field of palaeogenomics. By making molecular tip dating more accessible and reproducible, our pipeline will allow researchers to make their results more comparable across different datasets and taxa. From a more practical perspective, ancient DNA research groups can benefit from a quick and easy way to estimate the age of their samples, not only to place them in a better context within a project, but also to identify which specimens are most promising to target for future research.

## Contributions

Conceptualisation and Methodology: PDH, LD, JCCD

Pipeline development: WL, BS, JCCD

Pipeline testing and review: BS, WL

Pipeline validation: BS, WL

Funding acquisition: LD, JCCD

Project administration: JCCD

Supervision: JCCD

Visualisation: BS, WL, JCCD

Writing – original draft: BS, WL, JCCD

Writing – review & editing: All authors

## Acknowledgements

We are grateful to Edana Lord for her advice in bioinformatic analysis and to the staff at the Centre for Palaeogenetics (CPG) for their valuable discussions and beta testing during the development of the pipeline. The development of the pipeline was enabled by computational resources provided by the National Academic Infrastructure for Supercomputing in Sweden (NAISS), partially funded by the Swedish Research Council through grant agreement no. 2022-06725 (project ids: NAISS 2025/22-950 and NAISS 2025/22-1155).

## Data availability

The source code, documentation, and example datasets are available on GitHub at https://github.com/CpgSthlm/MitoDate.

